# BUDDY: An interactive web platform to accelerate the functional modeling of human missense variants using *Saccharomyces cerevisiae* orthologs

**DOI:** 10.64898/2026.10.19.760001

**Authors:** Maisee Brown, Aditya Sriram, Jeffrey Garcia, Brian M. Wasko

## Abstract

The advent of next-generation sequencing has transformed clinical genetics, accelerating the identification of human genetic variants. However, the labor-intensive process of functional validation of Variants of Uncertain Significance (VUS) remains a significant diagnostic barrier. While expressing humanized genes in *Saccharomyces cerevisiae* is a common approach for modeling missense variants with functional assays, using native yeast orthologs provides a more streamlined approach that preserves species-specific interactions within multisubunit complexes. However, taking this approach relies on a fragmented workflow that requires bioinformatic experience.

To address this issue, we developed the Bioinformatic Utility for Diagnostic Discovery in Yeast (BUDDY), an integrated web platform designed to streamline variant analysis by connecting bioinformatic tools and databases including ClinVar. Features include amino acid alignments, a protein structure viewer that displays a superposition of yeast and human proteins, pathogenicity prediction data, CRISPR-Cas9 reagent design, and yeast phenotypes to provide potential functional assays. The platform also contains an artificial intelligence (AI)-assisted experimental design feature that evaluates the utility of modeling specific variant and gene combinations in yeast and can assist with phenotype selection.

BUDDY significantly accelerated the bioinformatic preparation time needed for ortholog variant modeling by an experienced biologist. Additionally, without prior bioinformatics training, a first-year medical student used BUDDY to develop a self-directed project and proposed a study of five Lynch syndrome-associated VUS using a forward mutation rate assay.

BUDDY is designed to accelerate the investigation of human variants in yeast and has a high potential utility for use in resource-constrained settings such as course-based undergraduate research experiences (CUREs).

## Introduction

The use of next-generation sequencing in clinical genetics has accelerated the identification of human genetic variations, but it has created a diagnostic backlog of Variants of Uncertain Significance (VUS). Modern high-throughput methods such as deep mutational scanning (DMS) can evaluate many thousands of variants simultaneously (Fowler and Fields 2014), but are used to broadly map the variant landscape rather than more deeply exploring the molecular mechanism of individual variants. Additionally, DMS requires labor-intensive library preparation, high-throughput sequencing, and advanced bioinformatic analysis to distinguish biological signal from noise. In contrast, CRISPR-mediated modeling of individual target variants provides a more streamlined and accessible methodology, affording mechanistic understanding in conjunction with rapid experimental validation. By leveraging yeast models, the field can continue to provide clinical-translational data that can support improved diagnostics and precision medicine.

*Saccharomyces cerevisiae* (yeast) offers an ideal platform for modeling conserved variants, as many fundamental biological processes are highly conserved. Yeast also have a propensity for homology-directed repair and have exceptional genetic tractability. By directly modifying the native yeast genome, researchers can bypass the technical complexities associated with humanizing entire genes and expressing heterologous proteins, thereby significantly reducing the workload for functional validation. By directly introducing clinical mutations into endogenous yeast orthologs, researchers can quickly generate robust phenotypic readouts of variant effects. As a result, this streamlined yeast-based approach provides a scalable framework for systematically elucidating sequence-to-function relationships.

The decentralized bioinformatic pipeline required to identify and model variants in native yeast orthologs can still pose a technical hurdle, thereby reducing accessibility. A standard pipeline requires querying multiple databases such as ClinVar to identify variants to study, and using tools such as DIOPT (DRSC Integrative Ortholog Prediction Tool) to identify human and yeast gene orthologs. Next, amino acid sequence alignments are needed to map conserved human variants to their yeast equivalents. Additionally, 3D structures of the human and yeast proteins can be compared to evaluate the overall structure similarities as well as the structural conservation surrounding the variant of interest. One way to assess whether a given variant results in a functional consequence is to identify if it results in an established phenotype associated with loss-of-function mutations via a search of Alliance of Genome Resources (The Alliance of Genome Resources Consortium 2024) and/or the scientific literature. The researcher must then assimilate the collected information and make a judgment on whether the gene, variant, and phenotype make a feasible experimental design given their particular laboratory constraints. Then there remains the need for an understanding of CRISPR sufficient to design the synthetic DNA needed for the CRISPR system used. Collectively, this decentralized multi-step process is time-consuming and can act as a significant barrier to novice researchers such as students.

To accelerate the process for identifying human variants that are functionally assessable in yeast orthologs, we developed BUDDY (Bioinformatic Utility for Diagnostic Discovery in Yeast), a web platform designed to streamline the decentralized workflows (Fig. 1). By bringing together clinical databases, orthology mapping, pathogenicity predictions, and 3D structural modeling, BUDDY automates the human-to-yeast ortholog experimental pipeline for variant discovery and provides metrics to help evaluate the suitability of a yeast ortholog model. Furthermore, BUDDY deploys a multimodal Large Language Model (LLM), which accesses the retrieved multi-dimensional data to help guide the selection of a phenotypic readout and to evaluate the feasibility of modeling specific human variant–gene combinations in the orthologous yeast gene.

**Fig. 1.**
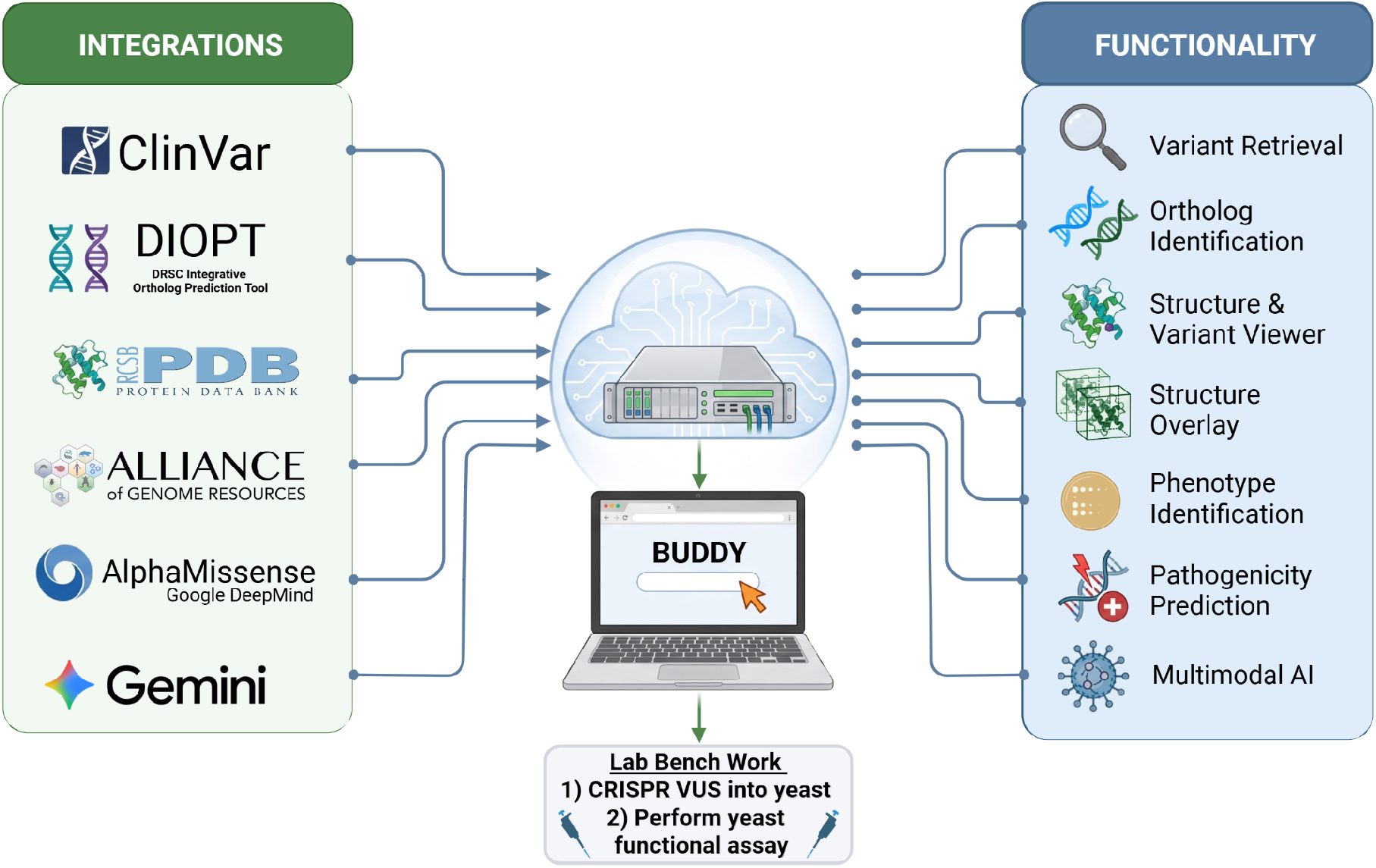
An overview of BUDDY’s data integrations and functionality. Created in BioRender. Wasko, B. (2026) https://BioRender.com/b5×693g.

BUDDY is designed to reduce the technical barriers to entry, making bioinformatic analysis and VUS modeling in yeast orthologs more accessible. This accessibility allows BUDDY to act as a potential educational framework, including for use within Course-based Undergraduate Research Experiences (CUREs) by connecting a safe, cost-effective, fast-growing model organism to uncharacterized variants in clinical genetics. BUDDY is meant to empower researchers and students to actively mitigate the growing pool of human VUS and expand the mechanistic understanding of disease.

## Materials and Methods

### Architecture and Implementation

BUDDY is a comprehensive web-based platform, architected as a full-stack application to facilitate conserved variant identification and experimental design. The client-side interface is constructed using React 19 and TypeScript, with Tailwind CSS providing the user experience. The server-side backend is powered by a Node.js/Express framework, which interfaces with external APIs to retrieve and integrate relevant data.

### API Dependencies

Our bioinformatic software integrates with various external APIs to retrieve and analyze biological data. REST and GraphQL APIs are used from prominent databases, including ClinVar (Landrum et al. 2025), the Alliance of Genome Resources (The Alliance of Genome Resources Consortium 2024), AllianceMine, MyVariant.info, MyGene.info (Xin et al. 2016), UniProtKB (UniProt Consortium 2025), AlphaFold Protein Structure Database, RCSB Protein Data Bank (PDB) and PDBe Graph API (Vallat et al. 2026), BioGRID (Oughtred et al. 2021), NCBI Entrez E-Utilities, and gnomAD (Karczewski et al. 2020). These APIs provide access to a wide range of data, including curated loss-of-function phenotypes, genomic DNA sequences, ClinVar classifications, protein structures, and population allele frequencies. Additionally, we leverage the Google Gemini API (Gemini Team 2025) for searching via natural language gene discovery and experimental assay generation, enabling integration with Google Search.

### Data Acquisition and Ortholog Mapping

To map human clinical data to *S. cerevisiae*, BUDDY fetches orthology scores derived from DRSC Integrative Ortholog Prediction Tool (DIOPT, Hu et al. 2026). Human variants are retrieved from ClinVar and AlphaMissense scores from dbNSFP (Liu et al. 2020) via the MyVariant.info API, with validation and conflict resolution conducted via NCBI E-utilities API. Once a human-yeast ortholog pair is established, canonical protein sequences are retrieved from UniProtKB and aligned using a dynamic pairwise alignment (Needleman-Wunsch global by default, or optionally via Smith-Waterman local alignment). Residue conservation is determined using a BLOSUM62 (Henikoff and Henikoff 1992) scoring matrix to classify residues as identical or similar. Furthermore, yeast phenotypic data are retrieved from Alliance of Genome Resources (*Saccharomyces* Genome Database) (Engel et al. 2022).

### Structural Visualization and Superposition

Protein structure visualization is natively embedded within BUDDY using 3Dmol.js (Rego and Koes 2015). The system fetches experimentally determined or AlphaFold-predicted structures (Bertoni et al. 2026) via UniProt or RCSB. To promote evaluation of the spatial conservation of selected proteins and variants, BUDDY provides a superposition of the human and yeast structures (Kabsch 1976; Zhang and Skolnick 2005; Meng et al. 2006) and allows the user to make manual overlay adjustments. The viewer highlights the variant, providing optional rendering styles (e.g., cartoon, sphere, stick). UniProt is also accessed to provide annotated features (e.g., domains, catalytic sites, metal binding sites, post translational modification sites), as well as 3D interaction interfaces derived from PDBe, which can be overlaid on both the amino acid alignment and the 3D viewer.

### CRISPR-Cas9 Reagent Design

To enable efficient in vivo modeling of variants in yeast, BUDDY features an automated design module for CRISPR-mediated genome editing reagents. This module identifies variant-proximal Protospacer Adjacent Motifs (PAMs) with a default selection for the widely-used *Streptococcus pyogenes* Cas9 (SpCas9) enzyme, which recognizes the NGG motif (Mojica et al. 2009). Small guide RNA (sgRNA) efficiency is predicted using a computational implementation of the Doench Rule Set 1 (Doench et al. 2014), promoting optimal sgRNA design. Additionally, BUDDY generates oligonucleotides for CRISPR-mediated genome editing as described by Laughery et al. (2015), including repair template oligonucleotides with homology around the sgRNA target site that introduce the desired amino acid substitution along with synonymous mutations in the PAM or seed region to prevent Cas9 re-cutting of the edited locus. Control genomic repair template oligonucleotides are also generated to introduce a frameshift loss-of-function (2-bp PAM deletion) mutation using the same sgRNA targeting plasmid. Furthermore, BUDDY can also design synthetic DNA sequences for an unpublished cloning-free (NoClo) yeast homology-directed recombination approach, utilizing plasmids from Bean et al. (2022). PCR primers flanking the CRISPR edit site are also generated for amplicon or Sanger sequencing to verify the genomic changes.

### AI-Assisted Experimental Design

An additional feature of BUDDY is its integration with a multimodal LLM, which enables the construction of a structured, multimodal prompt that leverages the collected data. This information sent to the multimodal model includes quantitative orthology metrics, DIOPT scores, and a local homology percentage (defined as the percent similarity between yeast and human residues within a 13-amino-acid window centered around the variant). The prompt also incorporates variant pathogenicity prediction data in the form of AlphaMissense scores (Cheng et al. 2023), loss-of-function phenotypes, and 2D visual representations of 3D structural superpositions between yeast and human proteins with the variant of interest indicated, as well as annotated features relevant at the selected residue position. Notably, the LLM is instructed to find relevant literature and citations through active web searches, utilizing Google’s Grounded Search to inform its suggestions. The prompt also allows users to define laboratory constraints, which by default are set to a “Classroom Safe” setting, prompting the LLM to filter out phenotypes that require highly toxic chemicals and instead select safer alternatives that are more suitable for a teaching laboratory environment. Furthermore, the prompt directs the LLM to search published DMS datasets, saturation mutagenesis studies, and MaveDB (Rubin et al. 2025) entries for the human and yeast genes to determine if comprehensive datasets regarding variants already exist.

### Estimates of Conserved Missense VUS from Human to Yeast

To evaluate the broad applicability of an ortholog based approach, we performed a genome-genome-wide variant comparison (which is not a feature of BUDDY). Ortholog pairs of yeast genes with a DIOPT (v10) score ≥2 were filtered to the top-scoring yeast target for each human gene. Canonical protein sequences for human (taxonomy ID 9606) and *S. cerevisiae* (taxonomy ID 4932) were retrieved using the UniProt REST API. Missense VUS from ClinVar (https://ftp.ncbi.nlm.nih.gov/pub/clinvar/tab_delimited/variant_summary.txt.gz released 2024-03-31 19:46:01) were filtered to exclude non-missense variants and then mapped to their corresponding orthologous genes. Using Biopython PairwiseAligner (v1.87; BLOSUM62, gap open: -5, extend: -0.5), variant positions were aligned to identify those located at conserved (identical or similar amino acids, BLOSUM62 score > 0) yeast residues. The conserved missense VUS were then quantified and stratified according to their DIOPT rank.

### Bioinformatic Processing Times

The human manual process encompassed retrieving yeast and human protein sequences, performing an amino acid sequence alignment, finding the position and conservation of the target residue, and designing oligonucleotide sequences for engineering the missense mutation and silent PAM mutation, as well as identifying a potential phenotype. A visual comparison of the 3D structures was omitted from the manual timing, which would have increased the duration for the human assessment. A paired t-test was performed to determine statistical significance.

## Results

### Estimating the Yeast Ortholog Tractable VUS Landscape

We performed an estimate of the potential landscape of VUS modelable using orthologous yeast genes. From 2,355,334 unique VUS in ClinVar, of which 2,128,197 were missense (90%), 313,562 missense VUS were identified that aligned at conserved residues of genes with a DIOPT score of >=2. Of those, 188,456 had a ‘high’ (reciprocal best match) DIOPT rank. This suggests that 9-15% of total missense VUS are potentially modelable in a yeast ortholog.

### Reduced Bioinformatic Processing Time

In user testing conducted by the corresponding author, an experienced yeast biologist, BUDDY significantly reduced the bioinformatic processing time from an average of 18.5 ± 8.7 min (n=4) to 0.96 ± 0.20 min (n=4, p=0.0271).

### BUDDY Output

The output of a sample gene, *ATP6V1B1*, is indicated in Fig. 2, displaying the amino acid alignment (Fig. 2a), phenotypes (Fig. 2b), the table of ClinVar variants (Fig. 2c), and the structure viewer (Fig. 2d). This information is meant to enable users to identify variants for modeling in yeast. Upon selecting a target, BUDDY can generate oligonucleotide sequences needed for genetic engineering (Fig. 3a). An optional AI-assisted experimental plan can also be generated, with a sample partial output including summary information indicated (Fig. 3b). The AI-assisted experimental plan selects from known yeast phenotypes associated with the gene to provide a potential assay to use for functional assessment of the variant.

**Fig. 2.**
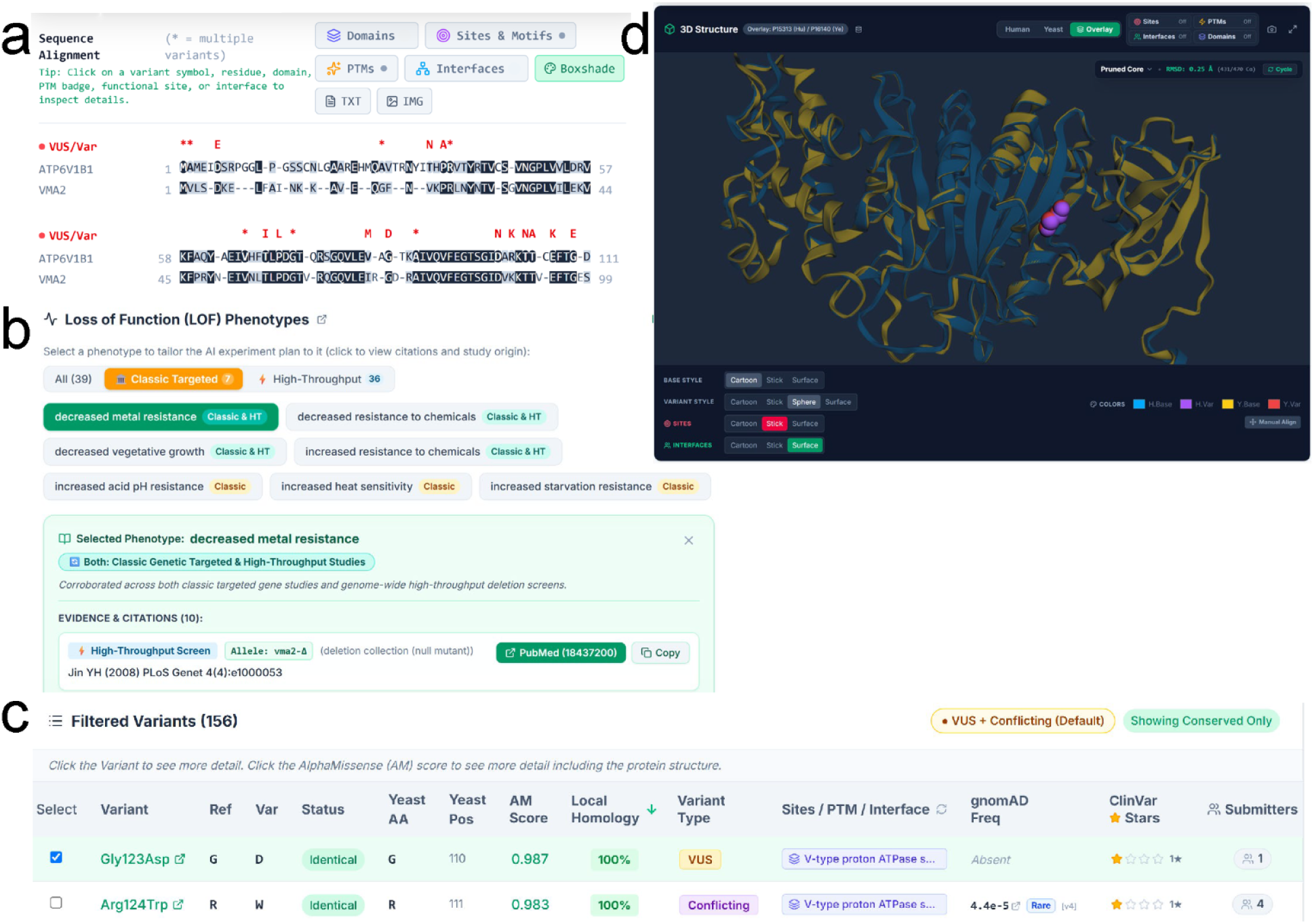
Sample output from BUDDY. Representative partial outputs from BUDDY displayed to users for *ATP6V1B1*, including the **a)** amino acid sequence alignment, **b)** phenotypes identified from Alliance of Genome Resources, **c)** table of variants, and **d)** structure viewer.

**Fig. 3.**
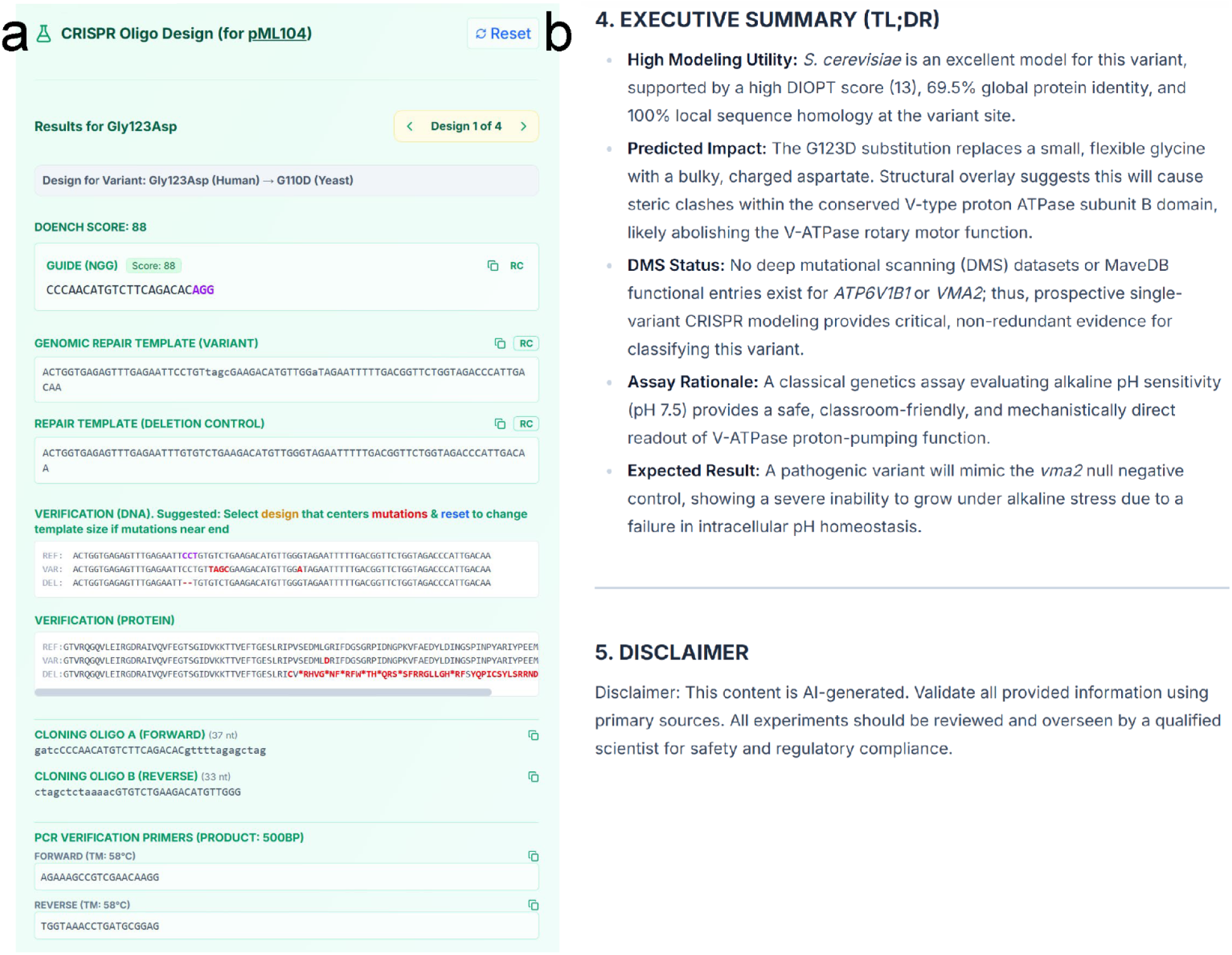
Additional sample output from BUDDY. Representative output for the **a)** CRISPR oligonucleotide design and **b)** partial output from the AI-assisted experimental design feature.

### Application of BUDDY

To evaluate the utility of BUDDY, a first-year medical student was tasked with using the software to investigate a self-selected gene related to their interests. Following a brief software demonstration, the student independently identified five VUS within conserved residues of their gene choice without prior bioinformatics training or further software support.

The gene *MSH2*, critical for DNA mismatch repair and driver of Lynch syndrome, was chosen for its relevance to cancer. The direct links to ClinVar submission information alongside the AlphaMissense prediction scores were utilized to choose the variants shown in Table 1. A *CAN1* canavanine resistance forward mutation assay (Whelan et al. 1979) was selected as a phenotype.

**Table 1.** Student-selected VUS with rationale.

| Variant | AlphaMissense Score | Rationale for Selection |
| --- | --- | --- |
| Lys449Arg | 0.151 | Selected to investigate the discrepancy between its high number of submissions (9) in ClinVar and yet a low AlphaMissense pathogenicity prediction score. |
| Glu455Gln | 0.348 | Chosen due to five independent submitter entries explicitly noting annotation of Lynch syndrome, yet a low AlphaMissense score. |
| Met672Arg | 0.984 | Selected due to conflicting entries (VUS and benign) within ClinVar while possessing a high AlphaMissense score. |
| Gly713Arg | 0.999 | Selected as a high probability hit due to a very high AlphaMissense score. |
| Thr724Met | 0.993 | Chosen due to conflicting ClinVar entries with multiple Uncertain significance entries as well as a Likely benign entry, yet a high AlphaMissense score. |

The use of BUDDY’s unified dashboard allowed variant retrieval, mapping to the yeast ortholog, and visualization of the human and yeast structures, a workflow that traditionally requires manually retrieving data across multiple independent sources. The student noted that different phenotypes were suggested by AI during deployment on different VUS (consistent with the stochastic nature of LLMs), and the mutational flux assay was ultimately selected as a phenotype for study.

Notably, this dry-lab platform testing led to the generation of multiple additional features within BUDDY, including a discordant search for variants with opposing AlphaMissense prediction scores and ClinVar submission classifications. Additionally, the LLM prompt was adjusted to specifically include grounded web searches for published deep mutational scanning (DMS) results related to the gene of interest, given the related DMS publications about *MSH2* variants (Jia et al. 2021; Ollodart et al. 2021).

## Discussion

BUDDY is designed to consolidate a fragmented, multi-source workflow to aid in modeling human VUS in orthologous yeast genes to allow rapid targeted functional assays. Additionally, BUDDY has the potential to serve as a scalable framework for use in CUREs, empowering students to investigate authentic clinical genetic questions without requiring advanced bioinformatic experience.

BUDDY has multiple limitations. Biologically, the use of *S. cerevisiae* as a model system inherently introduces constraints, as yeast lacks the complex tissue architecture, specialized signaling pathways, and post-translational modifications present in humans. This evolutionary divergence could lead to instances where human variants disrupting higher-order metazoan functions or requiring tissue-specific contexts fail to produce a scorable phenotype in yeast. Additionally, BUDDY’s current focus is on mutating endogenous yeast orthologs, which provides an approach that preserves native genomic context and native protein-protein interactions, but also limits the assessment to conserved residues. While using yeast orthologs reduces the number of modelable VUS, it also retains species-specific protein interactions that may be essential for protein subunits of large multimeric complexes.

A notable software limitation includes the reliance of BUDDY on external APIs for real-time data aggregation and continuous uptime and stability. Additionally, the integration of generative AI introduces the inherent risk of AI hallucinations, although this may be partially mitigated through the use of automated web-search grounding. Response variability is also an intrinsic property of LLMs. The AI outputs are strictly meant to serve as a hypothesis-generating support tool, and an embedded disclaimer prompts users to independently validate the output and retain mentor supervision.

To enhance the robustness and usability of BUDDY, future developments will focus on improving computational resilience and reducing wet-lab technical barriers. To minimize the impact of external API downtime, future software iterations will implement enhanced data caching and additional fallback querying mechanisms, reducing potential downtime. Additionally, we aim to develop a robust and highly accessible cloning-free CRISPR workflow in yeast, which would substantially reduce hands-on experimental time for genetic editing and thus increase accessibility for VUS characterization. By streamlining the bioinformatic processes, BUDDY can accelerate functional genomics for researchers, and allow for the democratization of VUS discovery in resource-constrained settings, including within teaching labs as CUREs.

## Software Availability

## Availability

BUDDY is currently available to use online at https://buddy.wasko.org. Gemini API access costs are currently fully covered, but future deployments may require the user to provide their own API key.

**Source Code:** https://github.com/Wasko-Lab/BUDDY/

**Archived Software Version (v2.2.1):** https://doi.org/10.5281/zenodo.23127633

**Software License:** The software is open-source via GNU Affero General Public License v3 (AGPLv3).

## Funding

This work was supported in part by the Google Cloud Research Credits program.

## Conflicts of interest

The authors have no relevant conflicts of interest to declare.

## Notes

### Competing Interest Statement

The authors have declared no competing interest.

